# Statins do not reduce the parasite burden during experimental *Trypanosoma cruzi* infection

**DOI:** 10.1101/2025.01.30.635783

**Authors:** Sarah Razzaq, Francisco Olmo, Suresh B. Lakshminarayana, Ying-Bo Chen, Shiromani Jayawardhana, Srinivasa P.S. Rao, John M. Kelly, Amanda Fortes Francisco

**Author notes:** Sarah Razzaq and Francisco Olmo contributed equally to this work. Author order was determined by drawing straws.

## Abstract

Cardiomyopathy is the most common pathology associated with *Trypanosoma cruzi* infection. Reports that statins have both cardioprotective and trypanocidal activity have generated interest in their potential as a therapeutic treatment. Using a highly-sensitive bioluminescent mouse model, we show that 5 days treatment with statins has no significant impact on parasite load. The free systemic concentrations fail to reach the level required for potency. Hence, clinical trials to investigate trypanocidal activity of statins lack experimental justification.

---

Chagas disease cardiomyopathy is the major clinical manifestation of long-term infection with the protozoan parasite *Trypanosoma cruzi*, and affects 20-30% of those infected. Pathology driven by persistent inflammatory responses result in a range of cardiac impairments, permanent structural changes in the myocardium, and increased mortality (1). The current consensus is that parasite persistence is necessary for the development of Chagas cardiomyopathy (2). However, the drugs currently available to treat *T. cruzi* infection have limitations in terms of efficacy, and toxic adverse effects can lead to early treatment termination (3). Crucially, benznidazole (BZ), the front-line therapeutic drug, did not reverse cardiac damage in a clinical trial (4). A drug that combines trypanocidal activity, with an ability to control host factors that mediate cardiac pathology, would be the holy grail of Chagas disease research.

Statins are a group of fungal metabolites that inhibit 3-hydroxy-3-methyl-glutaryl (HMG)-CoA reductase, the rate-limiting enzyme in cholesterol biosynthesis. In addition to cholesterol lowering activity, statins have anti-inflammatory and immunomodulatory properties. They also slow blood clotting, stabilize atherosclerotic plaques and can reduce cardiovascular disorders (5- 8). These diverse systemic effects have generated interest in exploring their potential for treating infectious disease (9). Statins are currently used by >200 million people to help lower the level of low-density lipoprotein cholesterol in the blood. Although some safety concerns have been raised, the overwhelming evidence suggests that the benefits of therapy far outweigh the risks (10). Currently, a proof-of-concept phase II clinical trial (11) is ongoing to determine if statins have a beneficial impact on inflammation and cardiac function in non-symptomatic chronically infected patients pre-treated with BZ or nifurtimox. Reports suggest that simvastatin can reduce both parasitaemia and cardiac parasite burden in an acute model of Chagas disease, as well as inducing anti-inflammatory responses (12). Lovastatin was also reported to be effective against *T. cruzi* epimastigotes, and to potentiate the therapeutic effects of the ergosterol biosynthesis inhibitor ketoconazole (13). In contrast, although simvastatin improved cardiac remodelling in *T. cruzi*-infected dogs, it was not effective at reducing circulating parasites (14). The aim of the current work was to assess different statins in a highly sensitive experimental model of Chagas disease and to investigate the extent of their *in vivo* trypanocidal activity.

First, we assessed the *in vitro* activity of fluvastatin (Lescol XL), pravastatin and simvastatin (purchased from Novartis Pharmaceuticals, EDM Millipore Corp. and Cayman Chemical Co., respectively) against the *T. cruzi* CL Brener strain (DTU VI). Each compound was considerably less effective than BZ in blocking the growth of extracellular epimastigotes (Table 1). Similarly, activity against intracellular amastigotes, the parasite life-cycle stage that replicates in the mammalian host, was greatly inferior to that of BZ. Simvastatin was the most potent statin tested against amastigotes (EC_50_ = 5.7 µM), but was still 10 times less effective than BZ.

**Table 1.**
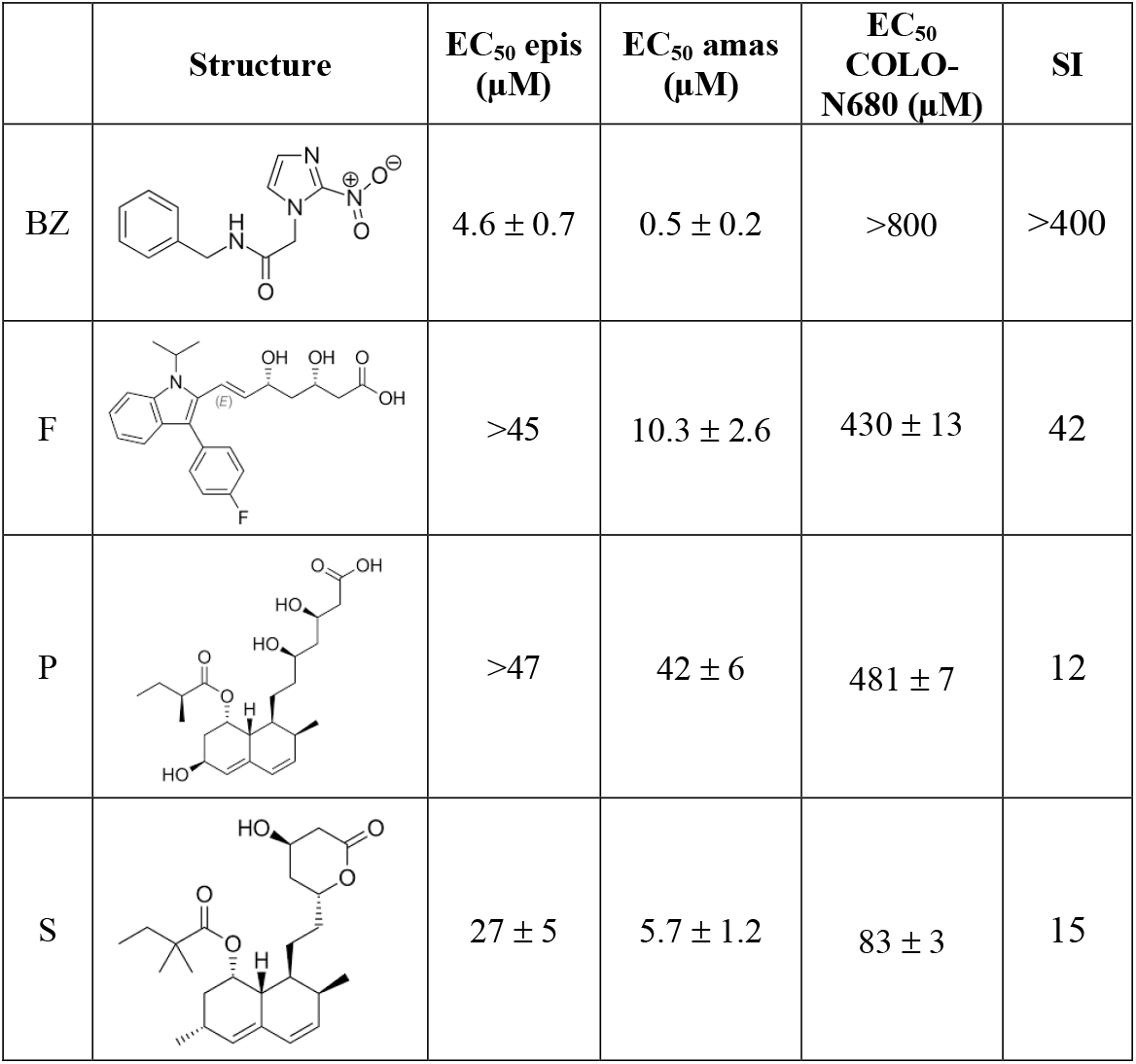
*In vitro* potency of statins against *T. cruzi* epimastigotes (epis) and amastigotes (amas). The activity of fluvastatin (F), pravastatin (P) and simvastatin (S) was assessed against the *T. cruzi* CL Brener Luc:mNeon strain (23) by applying eight-point potency curves (25). Benznidazole (BZ) was included as a standard. Mammalian cell cytotoxicity was determined using the COLO-N680 cell line. The selectivity index (SI) was the ratio of the amastigote/COLO-N680 EC_50_ values. Data was derived from two independent experiments carried out in triplicate (n = 6).

To assess *in vivo* efficacy, mice (aged 6 – 8 weeks) were infected with a strain of *T. cruzi* CL Brener engineered to express a bioluminescent fusion protein (15). At the peak of the acute stage, they were treated with 5 daily oral doses of fluvastatin, pravastatin and simvastatin, at levels that simulate daily exposure at the highest human doses (16) (Fig. 1A - D). None of the statin treatment schedules had any significant effect on the bioluminescence-inferred parasite burden, or the parasite organ/tissue distribution post-treatment. In contrast, BZ treatment (100 mg/kg) reduced the parasite burden by 99.8%, although by 35 dpi, parasite relapse was detected in each mouse (Fig. 1C). Mice typically require 20 days treatment with this BZ regimen to achieve sterile cure (17). Only simvastatin, the most active of the statins *in vitro* (Table 1), was tested as a treatment for chronic stage infection. We found that there was no significant impact on the parasite burden or organ/tissue distribution after 5-days treatment at 90 mg/kg, delivered 91 – 95 days post-infection (Fig. 1B - D). In contrast, treatment with BZ (100 mg/kg) reduced the parasite burden below the limit of detection. This BZ treatment schedule is generally curative when applied to chronic stage infections (9,10). At the experimental end-points (100 dpi, acute stage treatment; 175 dpi, chronic stage treatment), infection foci were prominent in the GI tract and skin, and sporadic in other organs and tissues (Fig. 1D), a pattern of distribution similar to that in non-treated mice in this infection model (15,18).

**FIG 1.**
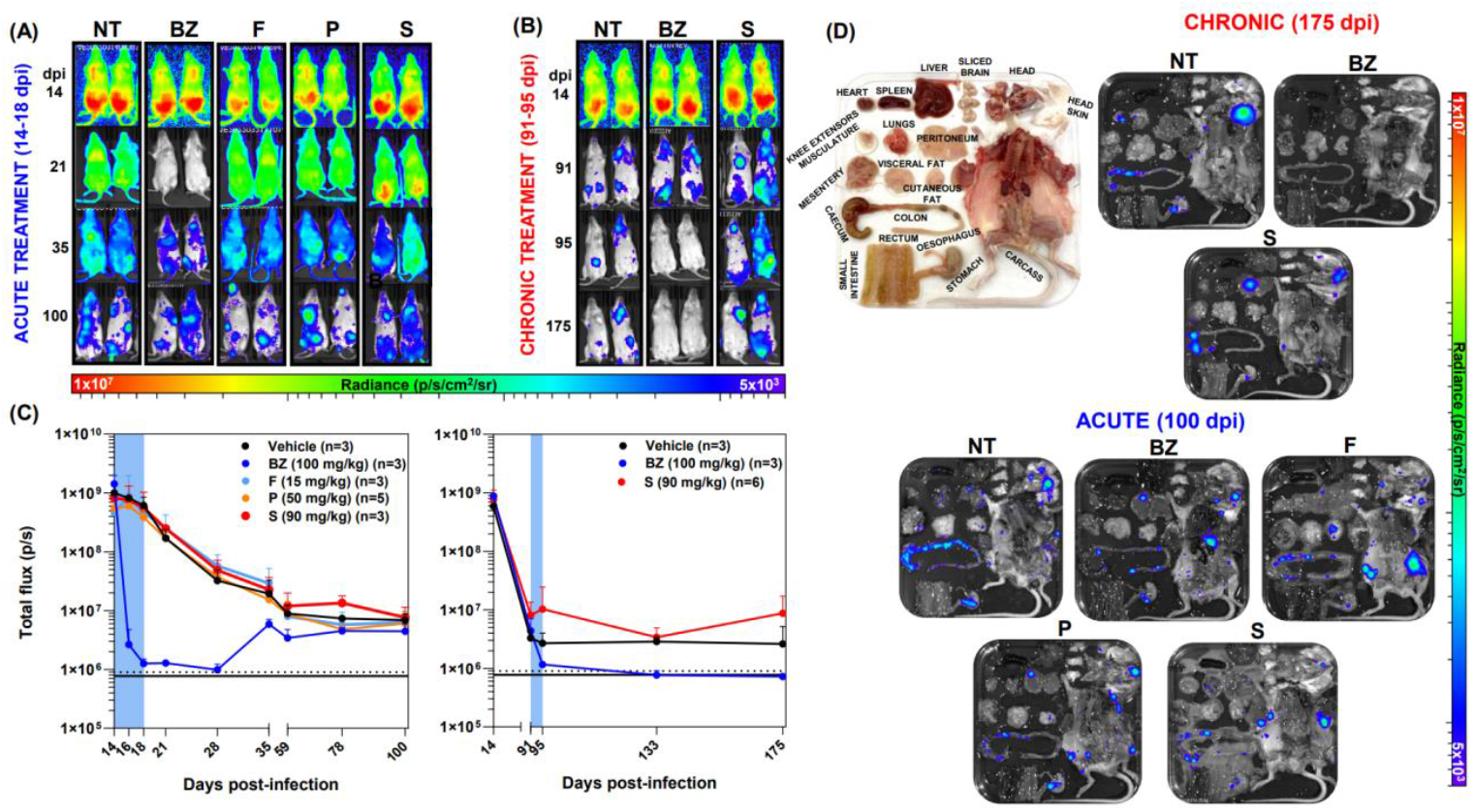
Statin treatment is ineffective at reducing the parasite burden during acute and chronic *T. cruzi* infections. (A-B) Representative *in vivo* ventral images of female BALB/c mice infected with 1x10^3^ bloodstream trypomastigotes of the *T. cruzi* CL Brener Luc strain (17). They were treated with benznidazole and statins for 5 days, beginning 14 days post-infection (dpi) for the acute treatment, and 91 dpi for the chronic treatment. Drugs were administered once daily by oral gavage. NT: non-treated (vehicle only); BZ: benznidazole-treated (100 mg/kg); F: fluvastatin-treated (15 mg/kg); P: pravastatin-treated (50 mg/kg); S: simvastatin-treated (90 mg/kg). The heat-map is on a log10 scale and indicates the intensity of bioluminescence from low (blue) to high (red); the minimum and maximum radiances for the pseudocolour scale are shown. (C) Graphs showing the mean bioluminescence (pixels/second; p/s) determined by *in vivo* imaging of treated and non-treated infected mice. Treatment groups (as above) and dosing regimens, including time of treatment (blue bar), are indicated. The black horizontal unbroken line indicates background bioluminescence established from non-infected mice (n=3), with the dashed line indicating SD above the average. (D) Representative *ex vivo* images of tissues and organs (24) from statin-treated mice (as above) during acute and chronic infections. Mouse *ex vivo* tissue/organ arrangement is shown in the picture display. Animal experiments were performed under UK Home Office project license P9AEE04E4 and approved by the LSHTM Animal Welfare and Ethical Review Board. All procedures were conducted in accordance with the UK Animals (Scientific Procedures) Act 1986.

In parallel, fluvastatin, pravastatin and simvastatin exposure were assessed in mice during acute stage infection (Fig. 2). Free statin concentrations were calculated based on their respective plasma protein binding. This revealed that the unbound levels of all statins remained below the concentrations required for *in vitro* amastigote potency (Table 1) for the duration of the testing period. In contrast, the unbound BZ concentration was maintained well above the amastigote EC_50_ value throughout (Fig. 2). Furthermore, other statin pharmacokinetic parameters predictive of bioavailability and *in vivo* efficacy were inferior to those of BZ (Table 2).

**FIG 2.**
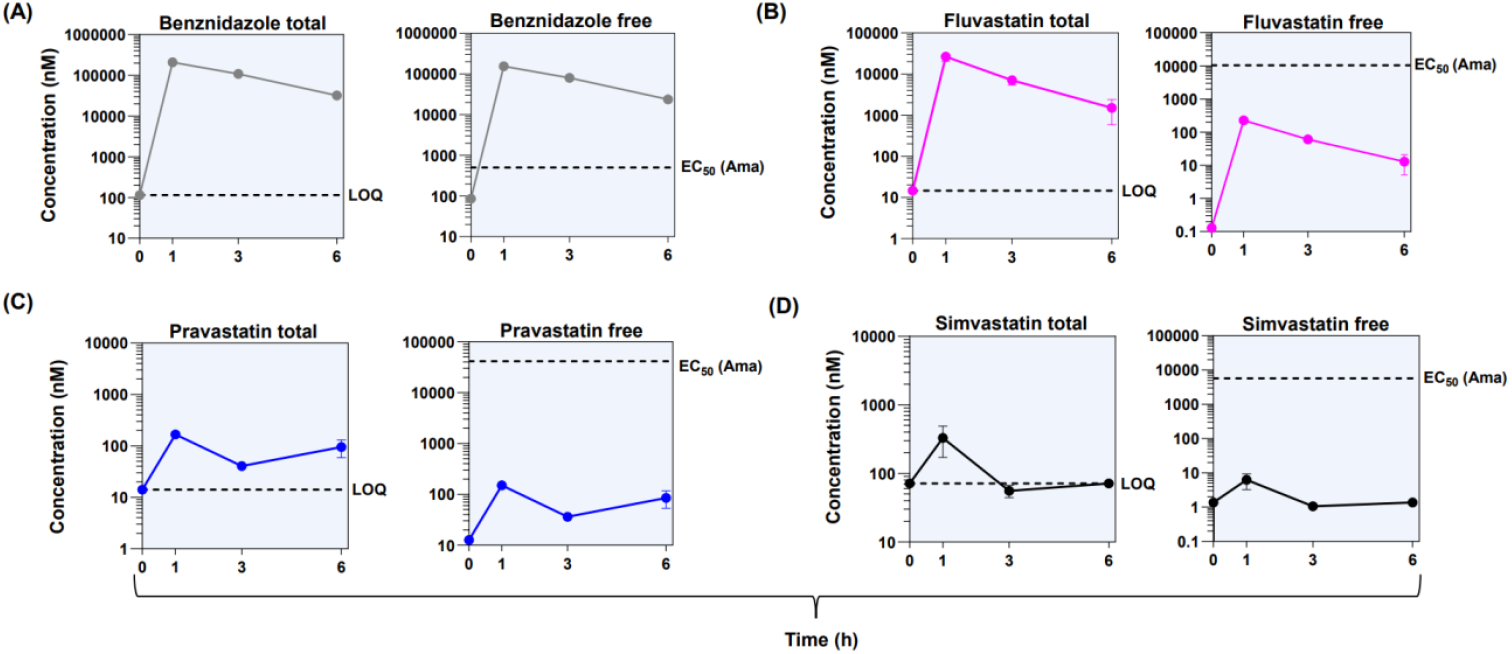
Systemic concentrations of statins and benznidazole during treatment of infected BALB/c mice. Drugs were administered by oral gavage at the doses described in the legend to Fig. 1. Following the last dose of acute stage treatment (day 18), blood samples were taken from the tail vein at 0 (pre-dose), 1, 3 and 6 hours, placed into cryovials containing 20 μl of milli-Q water, and stored at -20°C until analysis. Samples were prepared and analyte quantitation performed by optimized high-performance liquid chromatography coupled with tandem mass spectrometry (LC-MS/MS). Left-hand panels show the total systemic concentrations in each case, and which represent the average values from 3 mice. The dotted line represents the limit of quantification (LOQ). (A) benznidazole, 115 nM; (B) fluvastatin, 14.6 nM; (C) pravastatin, 14.1 nM; (D) simvastatin, 71.7 nM. The right-hand panels show mean free concentration ± SD, based on their respective plasma protein binding. The dotted line identifies EC_50_ against amastigotes (Table 1).

**Table 2.**
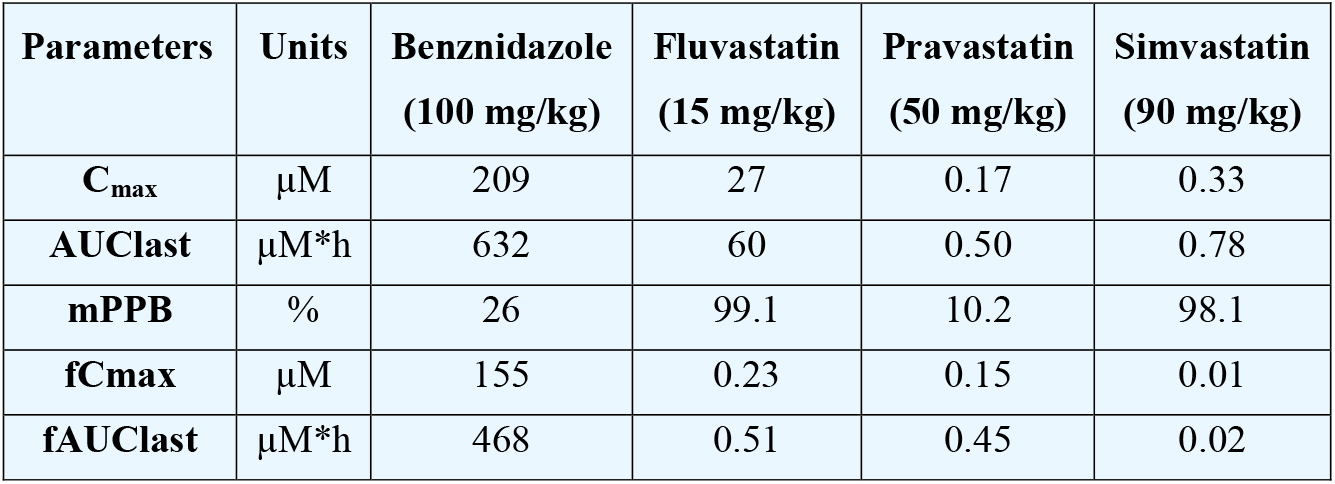
Pharmacokinetic parameters of statins in *T. cruzi* infected mice. Values were obtained using blood samples taken from treated female BALB/c mice (n=3) 18 days post-infection using the doses indicated (Fig. 2). Data from benznidazole-treated mice are shown for comparison. Cmax, maximum systemic concentration; AUClast, area under the curve, 0 – 6 hours; mPBB, mouse plasma protein binding; fCmax, maximum unbound systemic concentration; fAUClast; unbound area under the curve.

Several studies have reported that statins have potential for mitigating the development of chronic chagas heart disease (19-21), and a clinical trial to address this is underway (11). In addition, it has been suggested that statins may have an additional benefit, conferred through their trypanocidal activity (12,13,22). However, the available data on *in vivo* efficacy has been contradictory. Here, using highly-sensitive *in vivo* imaging employing a widely used murine model (23,24), we demonstrate that statins have no significant impact on the parasite burden during both acute and chronic *T. cruzi* infections. Consistent with this, we show that statin bioavailability is insufficient to produce an anti-parasitic effect. Therefore, when designing clinical trials to assess the therapeutic potential of statins against Chagas disease, their use in combination with other trypanocidal drugs as adjunct therapies (as in reference 11), represents the best evidence-based approach.

## AUTHOR CONTRIBUTIONS

Conceptualization, A.F.F.; methodology, F.O., Y.C., S.B.L. and A.F.F; software, S.R., F.O., Y.C., S.B.L. and A.F.F.; validation, S.R., F.O., S.B.L., Y.C.,S.P.S.R. and A.F.F.; formal analysis, S.R., F.O., S.B.L., Y.C., S.P.S.R. and A.F.F.; investigation, S.R., F.O., S.B.L., S.J. and A.F.F.; data curation, F.O., S.B.L., S.P.S.R. and A.F.F.; writing - original draft preparation, A.F.F.; writing - review and editing, F.O., S.B.L., S.P.S.R., Y.C., J.M.K. and A.F.F.; supervision, A.F.F. and J.M.K.; project administration, J.M.K.; funding acquisition, J.M.K. and A.F.F. All authors have read and agreed to the published version of the manuscript.

## ACKNOWLEDGMENTS

This research was supported by the UK Medical Research Council grants MR/T015969/1 to J.M.K. We would like to thank the Biological Services Facility team at LSHTM, especially James Gates and Carmen Abela for training, technical support, and scientific advice. We thank Linda Xiao and Colin Osborn from Novartis for their technical support and scientific advice. We declare no conflicts of interest.

